# Creating a genetic toolbox for the carbon-fixing, nitrogen-fixing and dehalogenating bacterium *Xanthobacter autotrophicus*

**DOI:** 10.1101/2024.06.26.600795

**Authors:** Alexa F. Van Voorhis, Rebecca S. Sherbo

## Abstract

*X. autotrophicus* is a metabolically flexible microorganism with two key features: 1) The organism has adapted to grow on a wide variety of carbon sources including CO_2_, methanol, formate, propylene, haloalkanes and haloacids; and 2) *X. autotrophicus* was the first chemoautotroph identified that could also simultaneously fix N_2_, meaning the organism can utilize CO_2_, N_2_, and H_2_ for growth. This metabolic flexibility has enabled use of *X. autotrophicus* for gas fixation, the creation of fertilizers and foods from gases, and the dehalogenation of environmental contaminants. Despite the wide variety of applications that have already been demonstrated for this organism, there are few genetic tools available to explore and exploit its metabolism. Here, we report a genetic toolbox for use in *X. autotrophicus*. We first identified suitable origins of replication and quantified their copy number, and identified antibiotic resistance cassettes that could be used as selectable markers. We then tested several constitutive and inducible promoters and terminators and quantified their promoter strengths and termination efficiencies. Finally, we demonstrated that gene expression tools remain effective under both autotrophic and dehalogenative metabolic conditions to show that these tools can be used in the environments that make *X. autotrophicus* unique. Our extensive characterization of these tools in *X. autotrophicus* will enable genetic and metabolic engineering to optimize production of fertilizers and foods from gases, and enable bioremediation of halogenated environmental contaminants.

## Introduction

Microbes have evolved to possess a wide array of metabolic processes based on their environmental conditions. We can harness these metabolic pathways to enable the sustainable utilization and conversion of waste and abundant resources into valuable products.^1^ For example, chemoautotrophs utilize CO_2_ and H_2_ as carbon and energy sources respectively, and these organisms can be metabolically engineered to produce liquid biofuels from gases.^2–5^ Nitrogen-fixing organisms can fix N_2_ into NH_3_ which can subsequently be used as a biofertilizer for plant growth^6–10^ in place of the energy-intensive Haber-Bosch process.^11^ In addition to gas conversion into fuels and fertilizers, some organisms also possess metabolic pathways that can decompose environmental contaminants such as halogenated compounds.^12–14^

Some species of microbes have evolved to host numerous beneficial metabolic pathways simultaneously due to habitat variation and environmental resistance. These species, known as habitat generalists, ensure survival in many environments by adapting to metabolize a broad range of small molecules.^15,16^ *Xanthobacter autotrophicus* is one key example: this gram-negative bacterium is a chemoautotroph, growing on CO_2_ as a carbon source and H_2_ as an energy source (Fig. 1a). *X. autotrophicus* can also use methanol, formate, and formaldehyde as carbon sources, oxidizing them with dehydrogenase enzymes to produce CO_2_ for the Calvin cycle.^17–19^ The dehydrogenase enzymes simultaneously convert NAD^+^ to NADH, providing energy for the cell. Furthermore, the strain *X. autotrophicus* Py2 can utilize propylene as a carbon source by performing an epoxidation reaction,^20–24^ while the strain *X. autotrophicus* GJ10 can dechlorinate dichloroethane and other aliphatic halogenated alkanes, alcohols and acids.^13,14^ These metabolic processes have been of great interest for removing halogenated environmental contaminants.^13,14^

**Figure 1.**
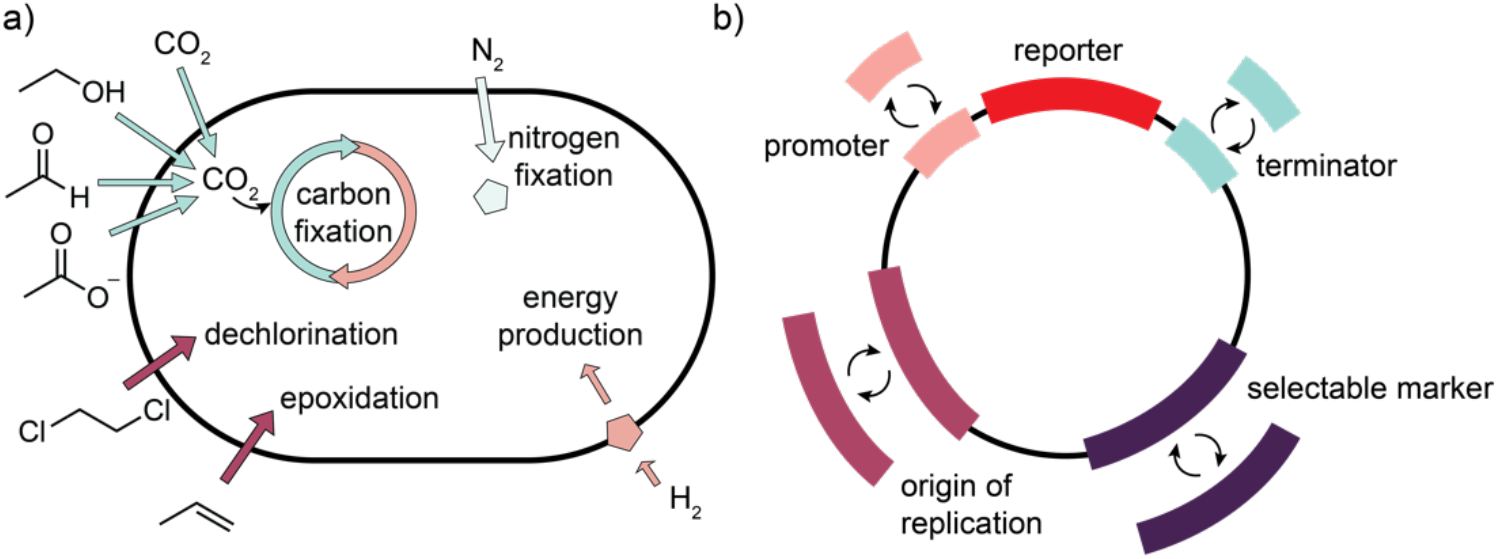
a) *X. autotrophicus* can derive carbon from CO_2_, methanol, formaldehyde, formate, chlorinated alkanes and acids, and propylene, nitrogen from N_2_, and energy from H_2_. b) We have identified a toolbox of genetic parts including origins of replication, selectable markers, constitutive and inducible promoters, and terminators that can be used to probe and engineer this organism’s unique metabolic processes, and we have demonstrated that these tools can be used under autotrophic and dehalogenative growth conditions.

In addition to the exceptional ability of *X. autotrophicus* to utilize a wide variety of carbon sources, this organism was the first chemoautotroph discovered that could also perform nitrogen fixation, using N_2_ as a sole nitrogen source (Fig. 1a).^25–27^ Growth on CO_2_, N_2_, and H_2_ is a rare metabolic combination because the processes have conflicting oxygen demands. O_2_ is required as an electron acceptor in the electron transport chain to produce enough ATP to power simultaneous carbon and nitrogen fixation. Yet, this O_2_ also inhibits both the RuBisCO and nitrogenase enzymes.^28,29^ Understanding this metabolic combination may be the first step in compatibilizing carbon and nitrogen fixation in other microbes and plants. This metabolic combination can also be used for the sustainable production of fertilizers^6^ and foods.^30,31^ The strain *X. autotrophicus* Py2 was grown on a combination of CO_2_, N_2_, and electricity-produced H_2_, and the resulting biomass was applied as a living biofertilizer, resulting in a ∼130% increase in the root mass of radishes.^6^ *X. autotrophicus* has been isolated in bacterial communities grown on CO_2_, N_2_, and H_2_ with the goal of producing microbial biomass for single-cell food protein,^30,32^ and we recently demonstrated the synthesis of the vitamin riboflavin starting from these same gaseous substrates by overproducing enzymes in the riboflavin pathway.^31^

The unique metabolism of *X. autotrophicus* has inspired the development of some tools for genetic modification. DNA editing has been performed through chemical^13,22^ and transposon^33^ mutagenesis, and DNA uptake has primarily been performed through triparental mating with cosmids.^13,19,22^ An effective electrotransformation protocol has also been reported for the strain *X. autotrophicus* GJ10.^34^ We have previously demonstrated overexpression of both reporter and native metabolic pathway genes using constitutive promoters derived from the *X. autotrophicus* genome.^31^ Despite these advances, translating standard genetic parts from model organisms to non-model organisms remains challenging,^35^ and even organisms within the same class can respond differently to the same part,^36^ so toolboxes must be specifically adapted for the genus or species.^37–40^

Here, we expand the genetic engineering tools available for use in *X. autotrophicus* (Fig. 1b) to enable the further exploration of its broad and versatile metabolism. We have chosen to use the strain *X. autotrophicus* GJ10 to create and test this toolbox because there is a facile electrotransformation protocol for DNA uptake,^34^ and because this specific strain can perform dehalogenation reactions relevant for bioremediation.^13,14^ We have implemented various origins of replication and determined their copy number and we have tested a set of antibiotic resistance cassettes for use as selectable markers. We have determined the promoter activity of several constitutive and inducible promoters and the termination efficiency of commonly used natural and synthetic terminators. Finally, we have demonstrated that the gene expression strategies reported here can be utilized under both the autotrophic and dehalogenative conditions that make the metabolism of *X. autotrophicus* unique.

## Results & Discussion

### Origin of Replication and Copy Number

We have tested four origins of replication in *X. autotrophicus* and determined that three origins resulted in successful plasmid replication (Fig. 2). We chose broad-host-range origins that have been successfully used in α-proteobacteria: RK2 (IncP), pSa (IncW), RSF1010 (IncQ), and BBR1 (Fig. 2a).^36^ RSF1010 is the most flexible origin because all the necessary proteins for replication initiation are incorporated within the origin,^41^ but it is large and the plasmid can be difficult to manipulate. BBR1 and pSa are small and stable origins that are easier to work with but are less flexible.^42^ RK2 is an intermediate option – this origin is smaller than RSF1010 but can be used in a broad range of hosts because replication is initiated with different proteins and in a different order depending on the organism.^43^

**Figure 2.**
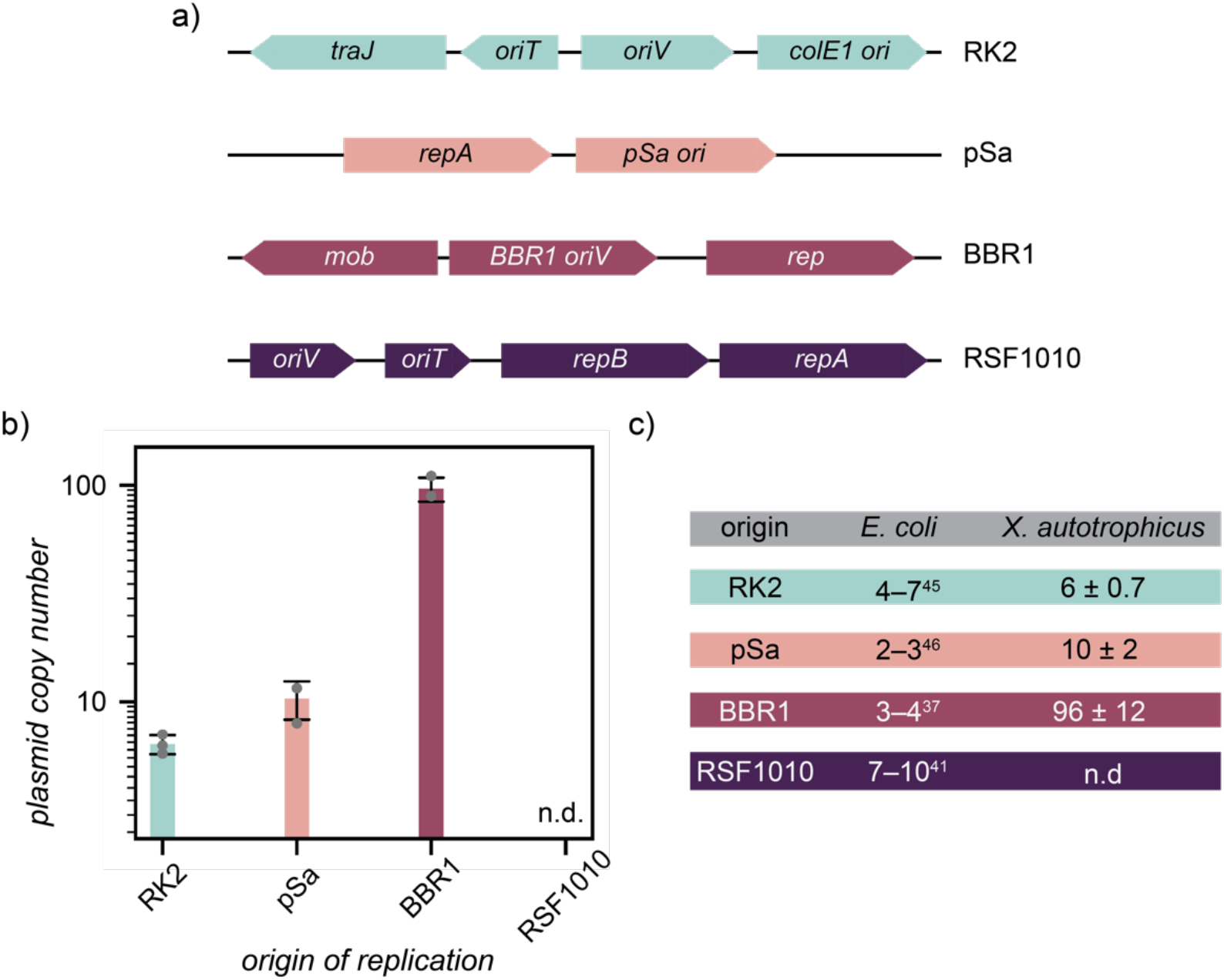
Origins of replication. a) RK2, pSa, BBR1 and RSF1010 origins of replication. b) Plasmid copy numbers for plasmids with three different origins of replication transformed into *X. autotrophicus*. RSF1010 did not result in successful electrotransformation to quantify copy number (n.d. = no data). c) Comparison between literature-reported plasmid copy number values for *E. coli*^37,41,45,46^ and the values reported here for *X. autotrophicus*.

We electrotransformed the four plasmids into *X. autotrophicus*, selected for transformants on antibiotic-containing plates, and successfully obtained colonies for every origin except RSF1010. Plasmids with the RSF1010 origin have also been unsuccessfully transformed in other α-proteobacteria,^36^ which may be the result of the metabolic burden that the extensive replication machinery imparts on the cell. We then utilized real-time PCR to determine the copy number for each successfully-transformed plasmid.^44^ We identified primer pairs that could selectively amplify the plasmid and chromosome and performed each amplification at three different DNA concentrations to measure qPCR primer efficiencies (Fig. S1). We used the cycle threshold (C_T_) and primer efficiency (E) for each strain to determine the copy number (see Materials and Methods). BBR1 resulted in ∼100 copies/cell, RK2 in ∼6 copies/cell and pSa in ∼10 copies/cell (Fig. 2b). Figure 2c shows the difference between literature plasmid copy number values in *E. coli*^37,41,45,46^ and the values obtained in this work for *X. autotrophicus*. There is good agreement for the low copy number origins RK2 and pSa. BBR1 has ∼30× higher copy number in *X. autotrophicus*, an effect that has also been observed for BBR1 in other organisms like *P. putida*.^37^ The existence of both low and medium copy number origins in *X. autotrophicus* provides options to balance gene expression and metabolic burden for a plasmid-based engineering approach.

### Selectable Markers

We have confirmed that kanamycin, gentamicin, ampicillin, and chloramphenicol are suitable selectable markers for *X. autotrophicus* (Fig. 3). Previous studies on antibiotic sensitivity in *Xanthobacter* suggested that the testing method and strain impacted whether the organism was sensitive or resistant.^17^ We have previously successfully used kanamycin as a selectable marker.^31^ To test new markers, we replaced *kanR* with *gmR, ampR*, and *cmR* genes, maintaining the same promoter in each case (Fig. 1b, selectable marker). We electrotransformed these plasmids into *X. autotrophicus* and plated on agar plates with three different concentrations for each antibiotic (Fig. S2), determining the appropriate concentrations as noted in Table S3. Each antibiotic resistant strain was grown in liquid culture along with the wild type and 20 μL aliquots of serial dilutions were plated on antibiotic-containing plates to demonstrate the difference between the strains (Fig. 3). In each case, single colonies could be isolated at 10^−4^−10^−5^ dilution factors and each culture contained 3×10^7^−2×10^8^ cfu mL^−1^ (Table S4), while no growth or colonies were observed at any concentration for the wild type strain. We have used *kanR* as a selectable marker for the remainder of this work.

**Figure 3.**
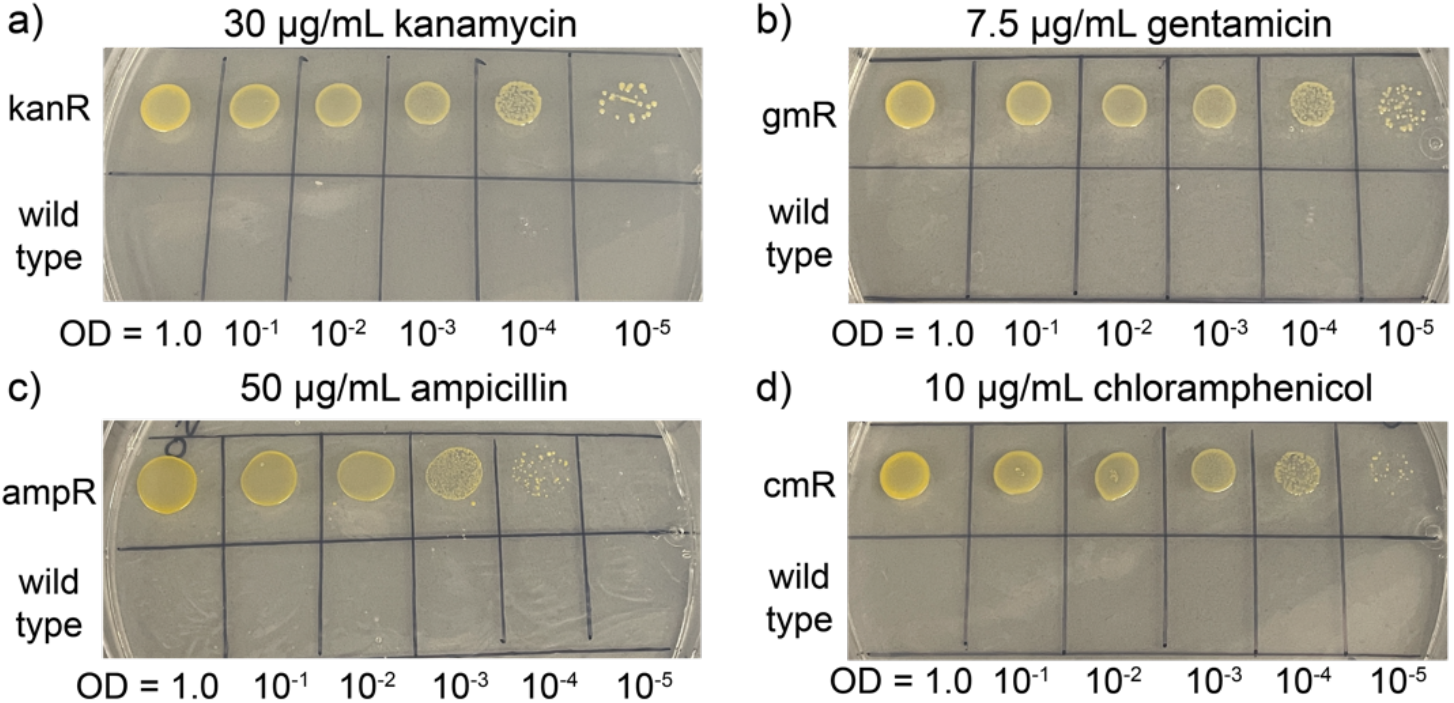
Antibiotic resistance markers. Modified strains containing plasmids with a) *kanR*, b) *gmR*, c) *ampR* and d) *cmR* antibiotic resistance genes grown on antibiotic-containing minimal media plates at a range of dilution factors compared to wild type *X. autotrophicus* GJ10. The wild type strain does not grow at any dilution factor.

### Promoters & Terminators

We tested six different constitutive promoters and achieved a range of gene expression levels in *X. autotrophicus* (Fig. 4). Five common constitutive promoters (pLacIq, pJ23103, pJ23107, pJ23102, pJ23111) were selected and compared against a strong constitutive promoter from the *X. autotrophicus* genome (pRpsM). We chose the pJ promoters from the Anderson series, a combinatorial library of varying-strength constitutive promoters,^47^ based on their high-, mid- and low-level activity in other nitrogen-fixing organisms.^7^ We placed a red fluorescent protein reporter gene (*mrfp1*) under control of this range of constitutive promoters (Fig. 4a) and performed plate reader experiments to measure red fluorescence intensity during growth. Figure 4b shows the growth-normalized fluorescence intensity for each promoter along with an RFP^−^ control. These promoters span a range of expression levels, with two promoters (pJ23102, pJ23111) having significantly higher *mrfp1* expression and resulting in visibly pink colonies (Fig. S3). These results are consistent with prior literature reports on other nitrogen-fixing organisms, where pJ23102 resulted in high expression levels in *Rhizobium* sp. IRBG74 and pJ23111 resulted in high expression levels in *P. protogens* Pf-5.^7^ The high expression levels resulting from pJ23102 and pJ23111 do impact growth densities and growth rates – these cultures reach peak optical density after ∼17 h instead of ∼14 h and grow to ∼0.1 optical density unit less than the other strains (Fig. S4). This growth impairment suggests that high fluorescent protein production results in a metabolic burden and, in cases where this burden is too high, the remaining promoters provide a range of expression levels all significantly higher than the RFP^−^ control.

**Figure 4.**
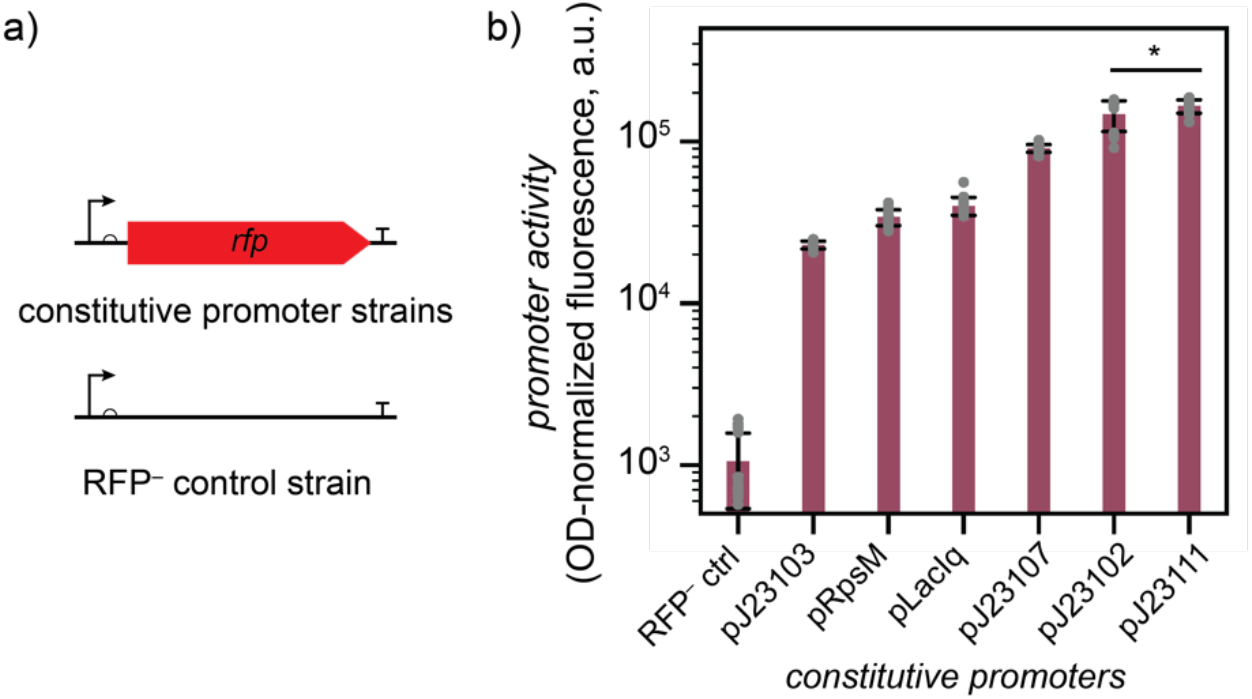
Constitutive promoters. a) Promoter activity was determined by using a red fluorescent protein reporter and keeping the ribosome binding site and terminator constant. An RFP^−^ strain was included as a control. b) Promoter activity of six constitutive promoters was measured by normalizing the red fluorescence (λ_ex_ = 577 nm; λ_em_ = 607 nm) to growth (optical density (OD) at 600 nm). The bars indicate the mean, the error bars indicate the standard deviation, and the grey dots indicate individual data points. A two-sample student t-test with unequal variances was performed to compare promoters with similar activities and * indicates P < 0.05.

We then tested four inducible promoters to determine whether an inducible promoter system could be successfully implemented in *X. autotrophicus* (Fig. 5). We chose pLac (lactose-inducible promoter with IPTG as an inducer), pTet (tetracycline-inducible promoter with anhydrotetracycline as an inducer), pCin (quorum sensing-derived inducible promoter with O3-C14 homoserine lactone as an inducer), and pBAD (arabinose-inducible promoter with arabinose as an inducer) for use in this organism. We first noted that each inducible promoter resulted in very little leaky expression, with the no-inducer controls resulting in similar fluorescence levels to an empty vector control (Fig. 5, Fig. S5). pLac and pTet resulted in low expression levels on induction, with a maximum 2× increase over the no-inducer control (Fig. 5a, b). We increased IPTG and aTc inducer concentrations to determine if expression level could be further optimized, particularly for pLac because a sigmoidal response curve was not observed, but increased inducer concentrations resulted in diminished cell growth and fluorescence intensity.

**Figure 5.**
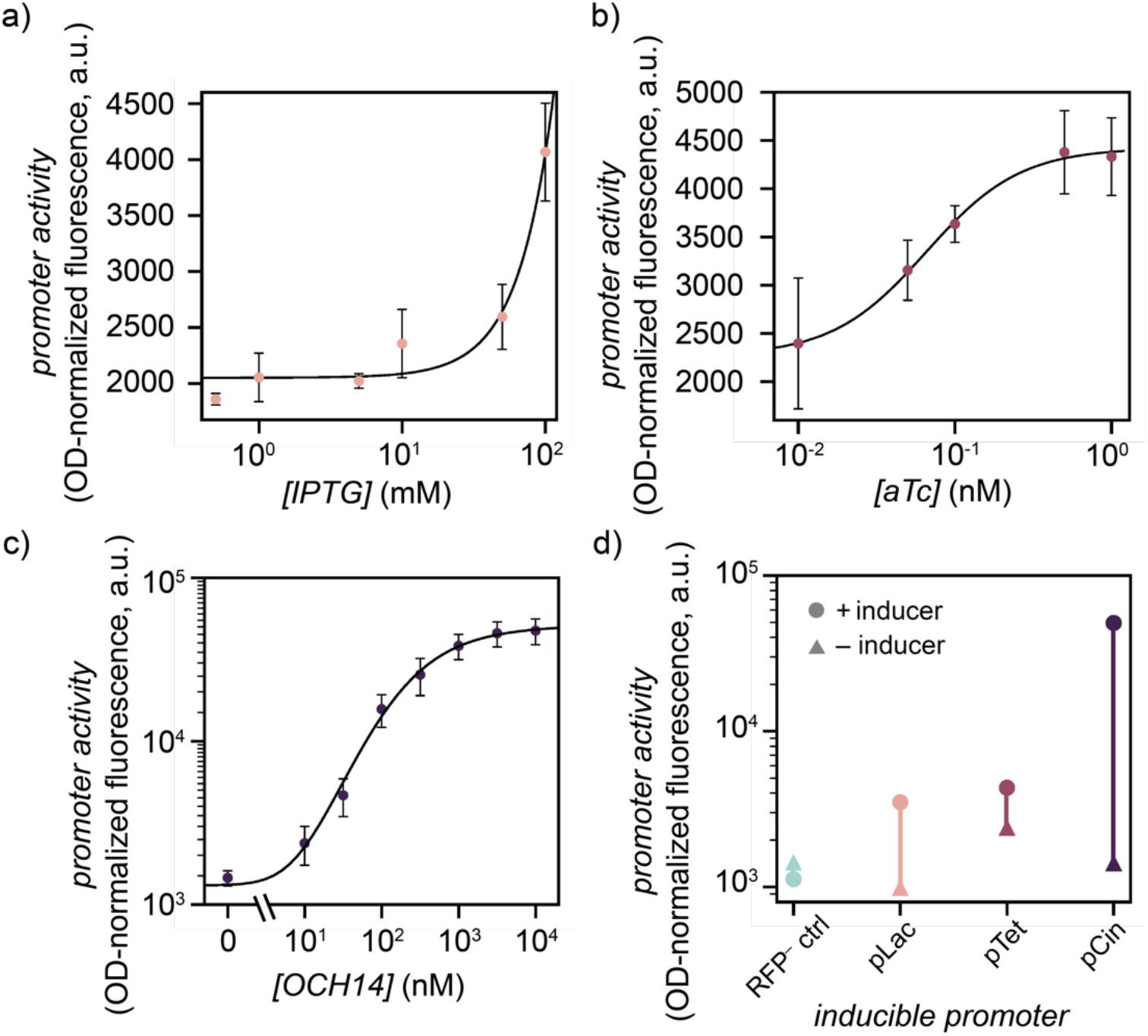
Inducible promoters. Promoter activity for the inducible promoters a) pLac/LacI with an IPTG inducer, b) pTet/TetR with an anhydrotetracycline (aTc) inducer and c) pCin/CinR with a 3-oxo-C14-homoserine lactone (OCH14) inducer. Promoter activity was measured at varying inducer concentrations by normalizing the red fluorescence (λ_ex_ = 577 nm; λ_em_ = 607 nm) to the optical density (OD) at 600 nm. The data points indicate the mean and the error bars indicate the standard deviation. Curves were fit using a symmetrical sigmoidal fit method based on the data points. d) Dynamic range for each inducible promoter compared to an RFP^−^ control. Triangles indicate promoter activity with no inducer and circles indicate promoter activity with the highest inducer concentration.

The pCin promoter resulted in a 25× increase in fluorescence intensity over 3 orders of magnitude increase in inducer concentration (Fig. 5c), demonstrating a large dynamic range and the ability to finely tune gene expression based on inducer concentration. The standard sigmoidal dose-response curve was observed, and the resulting fluorescence level at high inducer concentrations was similar to mid-level constitutive promoters like pLacIq. The final inducible promoter, pBAD, resulted in no fluorescence at any inducer concentration (Fig. S6a). We hypothesized that the pBAD promoter may not work in *X. auototrophicus* because the organism lacks arabinose import proteins – an issue that has been reported in other nitrogen fixers.^7^. To address this problem, we developed a new strain with a constitutively-expressed *araE* gene encoding an arabinose importer. Even with this modification, the pBAD strain still lacked a fluorescence response at any arabinose concentration (Fig. S6b), suggesting that pBAD is not effective inducible promoter system. Collectively, these results suggest that pCin has the largest dynamic range for induction of gene expression in *X. autotrophicus* (Fig. 5d).

We then examined a range of natural and synthetic terminators and determined that the synthetic terminators resulted in higher termination efficiencies (Fig. 6). We tested termination efficiency by placing the red fluorescent protein *mrfp1* and green fluorescent protein *gfp* genes under control of a single constitutive promoter and placing a terminator in between these reporter genes (Fig. 6a, Fig. S7). A GFP^−^ control had a terminator with no *gfp* gene and a terminator^−^ control had no terminator in between the fluorescent protein genes (Fig. 6a). Termination efficiency was determined by normalizing green GFP fluorescence to the GFP^−^ control, dividing by red RFP fluorescence, and normalizing this ratio to the terminator^−^ control (see Materials and Methods for details).^48,49^ This metric is a percentage that describes how often transcription stops vs. continues into the second fluorescent protein. Figure 6b shows that the strongest terminators (B1006, rrnB− T7TE double terminator) had termination efficiencies close to 100%, and there is no statistically significant difference between them. The three natural terminators had lower efficiencies which may impact growth and gene expression and are not optimal choices in strain construction.

**Figure 6.**
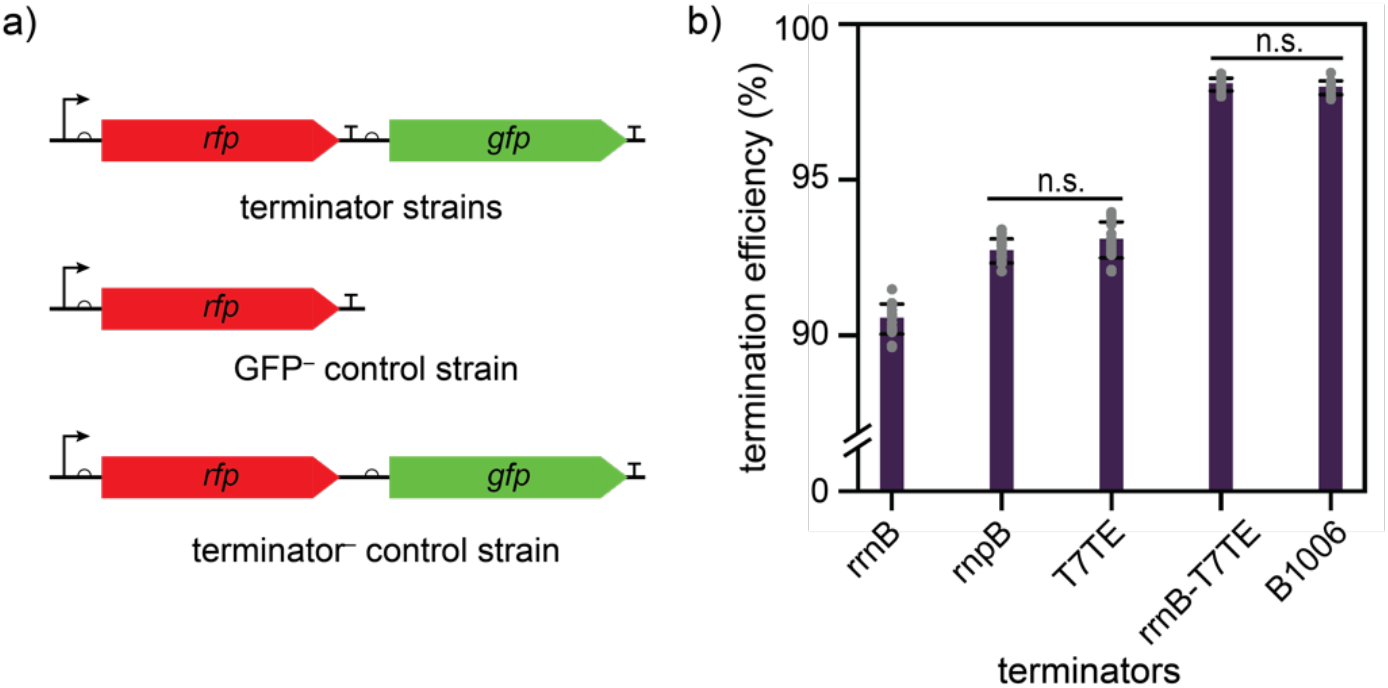
Terminators. a) Terminator activity was determined by placing a terminator between red and green fluorescent reporter genes. A GFP^−^ strain and a terminator^−^ strain were used for normalization. b) Termination efficiency was measured by dividing the green fluorescence (λ_ex_ = 485 nm; λ_em_ = 515 nm) by the red fluorescence (λ_ex_ = 577 nm; λ_em_ = 607 nm) and normalizing to the terminator^−^ control (see Materials and Methods). The bars indicate the mean termination efficiencies, the error bars indicate the standard deviation, and the grey dots indicate individual data points. A two-sample student t-test with unequal variances was performed to compare terminators with similar activities and n.s. indicates P > 0.05.

### Controlling the Metabolism

Finally, we demonstrated that we could control and maintain gene expression while growing *X. autotrophicus* under autotrophic conditions using methanol (MeOH) as a CO_2_ carbon source, and under dehalogenative conditions using dichloroethane (DCE) as a halogenated carbon source (Fig. 7). In this work, we have adapted commonly used promoters, terminators, origins of replication, and antibiotic resistance markers for use in *X. autotrophicus*. However, these parts must also function under the metabolic conditions that make this organism unique. We tested the MeOH and DCE carbon sources with two different promoters, pJ23111 (constitutive) and pCin (inducible), to confirm that we can control gene expression regardless of the carbon source. We utilized MeOH as a more facile, liquid autotrophic carbon source that provides CO_2_ to the Calvin cycle through a series of dehydrogenation steps.^18,19^ When DCE is used as a sole carbon source, a series of dehalogenases and dehydrogenases form glycolic acid, which then enters the central metabolism.^13,14^ In both cases, the carbon source is volatile so the culture tubes were sealed for growth. DCE is also only slightly miscible with water, so culture growth relied on shaking to ensure dispersion.

**Figure 7.**
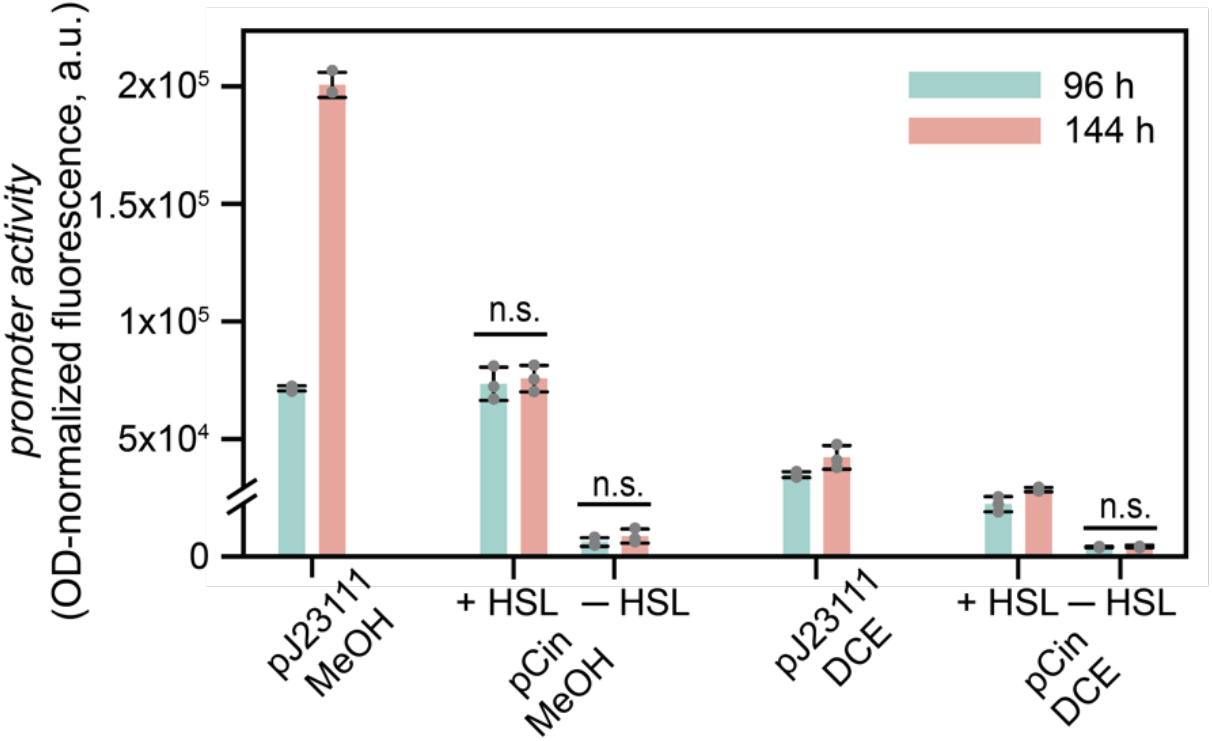
Promoter activity for the pJ23111 constitutive promoter and the pCin inducible promoter (+/– homoserine lactone inducer) with methanol (MeOH) or dichloroethane (DCE) as the carbon source. Measurements are taken at either 96 h (blue) or 144 h (pink). Promoter activity was measured by normalizing red fluorescence (λ_ex_ = 577 nm; λ_em_ = 607 nm) to the optical density (OD) at 600 nm. The bars indicate the mean, the error bars indicate the standard deviation, and the grey dots indicate individual data points. A two-sample student t-test with unequal variances was performed to compare conditions with similar promoter activities and n.s. indicates P > 0.05.

The constitutive promoter pJ23111 was active when grown on both MeOH and DCE, with higher expression levels in MeOH (Fig. 7). The inducible promoter pCin followed a similar trend, with improved expression in MeOH compared to DCE. The promoter activities for both pJ23111 and pCin were very similar when grown on MeOH as a carbon source compared to heterotrophic conditions. When no HSL inducer was added to the pCin cultures, promoter activity remained very low with both carbon sources, indicating very little leaky expression and consistent with previous heterotrophic experiments. Growth in both cases proceeded more slowly than with a standard heterotrophic carbon source, but expression continued at all timepoints sampled, suggesting that the slow growth does not impact our control mechanisms. These results confirm that the tools we present here can be applied in the environmental conditions that make *X. autotrophicus* unique.

## Conclusions

Here, we present a genetic toolbox for use in *X. autotrophicus*, an organism that can fix CO_2_ and N_2_ simultaneously and can use a range of C1 and chlorinated carbon sources. We have shown that both low and medium copy number broad-host-range origin plasmids can be utilized and have demonstrated and quantified resistance for four antibiotic resistance cassettes. We have identified constitutive promoters with a range of expression levels in this organism and have identified one high dynamic range and two low dynamic range inducible promoters. We have also quantified the termination efficiency of both synthetic and natural terminators. Finally, we grew strains with both a constitutive and an inducible promoter under carbon-fixing and dechlorination conditions and demonstrated that these genetic parts are still active and control expression under these conditions. We aim to provide tools to metabolically manipulate *X. autotrophicus* to enable further exploration and exploitation of its carbon- and nitrogen-fixing metabolism, its C1 carbon utilization, and its ability to dechlorinate environmental contaminants.

## Materials and Methods

### Organisms, Media and Reagents

*X. autotrophicus* GJ10 was received from the American Type Culture Collection (ATCC 43050) and re-streaked multiple times to isolate a slime-free mutant that was used here. *E. coli* DH5α competent cells were used for strain creation and were obtained from New England Biolabs (C2987H) or Invitrogen (18258012). *E. coli* was grown with LB media (10 g L^-1^ tryptone, 5 g L^-1^ yeast extract, 10 g L^-1^ NaCl) and *X. autotrophicus* was grown on minimal media with yeast extract and succinate (YESMM, Table S1). Antibiotics were added in the following concentrations as needed: kanamycin 50 µg mL^-1^ *E. coli*, 30 µg mL^-1^ *X. autotrophicus*; gentamicin 50 µg mL^-1^ *E. coli*, 20 µg mL^-1^ *X. autotrophicus*; ampicillin 100 µg mL^-1^ *E. coli*, 50 µg mL^-1^ *X. autotrophicus*; chloramphenicol 30 µg mL^-1^ *E. coli*, 10 µg mL^-1^ *X. autotrophicus*. All growth was performed at 37 °C and 220 rpm shaking.

All chemicals were used as received. All rich media components and agar were purchased from Difco. All minimal media components were purchased from Fisher Scientific. Inducers were purchased from Ambeed, VWR, and Fisher Scientific. All reagents for cloning and PCR were purchased from NEB and Fisher Scientific.

### Preparing X. autotrophicus Competent Cells

Two 100 mL cultures of *X. autotrophicus* grown in YESMM were grown to an OD = 1.0. Each culture was centrifuged at 13,000g for 10 min at 4 °C, washed with volume cold 1 mM NaOH, ½ volume cold dH_2_O, and ½ volume of cold 10% glycerol. The supernatant was removed, and the resulting cells were resuspended in 10 mL 10% glycerol and split up into 40 µL competent stocks.

### Strain Creation

Plasmids were assembled from the existing plasmids described in Table S2 by Gibson Assembly. Primers were designed by NEBuilder and DNA was amplified, digested with DpnI to remove original plasmid, and assembled with NEB HiFi DNA Assembly Master Mix. The resulting assemblies were transformed into *E. coli* DH5α competent cells. The assembled plasmids were isolated by miniprep and confirmed by whole-plasmid sequencing.

The plasmids were transformed into *X. autotrophicus* GJ10 by adding 2–4 µL plasmid to a 40 µL competent stock and incubated on ice for 10 min. The cells were added to a 2 mm electroporation cuvette and were electroporated at 2.5 kV, 25 µF and 600 Ω, resulting in a ∼15 ms exponential decay. The cells were recovered in 1 mL YESMM and incubated for 2 h before plating 100–400 µL on YESMM plates with the desired antibiotic concentration.

Colonies could be observed 2–3 days after plating and were re-struck on plates with the appropriate antibiotic. The presence of the correct plasmid was confirmed by colony PCR. A glycerol stock was made, and the glycerol stock was restruck on a YESMM plate with the correct antibiotics. The colonies of these re-struck plates were used to inoculate liquid media for the assays described below.

### qPCR and Copy Number Quantification

A 3 mL culture of each strain was grown for 48 h in YESMM, spun down, and DNA isolation was performed using a ZymoBIOMICS DNA kit with bead bashing. This isolation method was used because *X. autotrophicus* cells are difficult to lyse by traditional methods.^17^ Two primer sets were developed to perform qPCR: one that targeted the nitrogenase protein α-chain gene (132 bp, 80.2 °C T_m_ amplicon) and one that targeted the *mrfp1* gene in each plasmid (109 bp, 75.1 °C T_m_ amplicon). qPCR was performed with DNA diluted with 3 10-fold dilutions with a SYBR green master mix. qPCR was performed on a Bio-Rad CFX96 real-time PCR system with the following protocol: initial denaturation for 20 s at 95 ºC followed by 40× cycles of 3 s at 95 ºC and 30 s at 60 ºC. A final cycle of 15 s at 95 ºC, 60 s at 60 ºC, 15 s at 95 ºC and 15 s at 60 ºC completed amplification. The C_T_ values were plotted vs. DNA concentration to determine the primer efficiency. The plasmid copy number was then determined from the following formula:

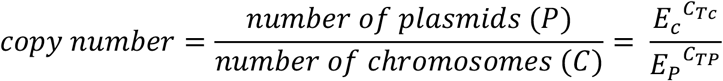

where E_C_ is the amplification efficiency of the chromosome, E_P_ is the amplification efficiency of the plasmid, C_TC_ is the cycle threshold for the chromosome, and C_TP_ is the cycle threshold for the plasmid. The copy numbers were determined from 3 replicates at 3 DNA concentrations for each strain and the error bars in Fig. 2 represent the standard deviation of these measurements.

### Antibiotic Resistance Assays

Electrotransformations were performed on the kanR, gmR, ampR, and cmR strains and 200 µL of cells in recovery media were plated on the following antibiotic concentrations: 10, 30 and 50 µg mL^-1^ kanamycin; 5, 7.5 and 10 µg mL^-1^ gentamycin; 50, 100 and 200 µg mL^-1^ ampicillin; and 5, 10 and 50 µg mL^-1^ chloramphenicol. The antibiotic concentration was decided by the colony count on each plate (Fig. S2). In each case, few and isolated colonies were observed, and all contained the plasmid, indicating that the antibiotic was effective against wild type cells. To further demonstrate this, 20 uL OD = 1.0 culture of each were plated on antibiotic plates along with 10-fold dilutions comparing the resistant and wild type cultures. The plates were incubated over 3 days and the wild type and resistant strains were compared (Fig. 3).

### Fluorescence Assays

#### Sample Preparation

One colony from the re-struck strain was transferred to 3 mL YESMM liquid media and grown for 48 h at 37 °C and 220 rpm. After 48 h, the culture was spun down and resuspended in 800 uL fresh YESMM. The optical density (OD) of the re-suspended culture was measured, and the cultures were diluted into a 96-well plate. Typically, ∼ 10 uL culture was added to 190 uL YESMM to make a total volume of 200 uL per well and a starting OD ∼ 0.1. Plate reader experiments were performed on a BioTek Synergy H1. The plates were incubated at 37 °C and orbital shaken at 237 rpm.

#### Constitutive Promoters

Cultures were grown in the plate reader for 17 h and both the OD_600_ (optical density at 600 nm) and mRFP1 fluorescence (λ_ex_ = 577 nm; λ_em_ = 607 nm; gain 125) were measured every 15 min. 6 replicates were performed for every strain on a single well plate (single colony), and 3 replicate plates were measured with a different colony for each plate, giving a total of 18 replicates. Data analysis was performed by identifying the peak in the growth curve for each strain and each replicate (*X. autotrophicus* growth curves typically decrease after a peak at the end of exponential phase due to a change in cell shape). The peak OD_600_ was averaged over 3 datapoints, and the corresponding fluorescence data was averaged at the same 3 timepoints. The fluorescence was normalized to the optical density. The average fluorescence for all replicates is plotted in Fig. 4 and the error bars correspond to the standard deviation. Statistical analysis was performed by using a two-sample student t-test with unequal variances and determining whether the P value was > or ≤ 0.05.

#### Inducible Promoters

Cultures were grown in the plate reader for 17 h and both the OD_600_ and mRFP1 fluorescence (λ_ex_ = 577 nm; λ_em_ = 607 nm; gain 125) were measured every 15 min. 2 replicates were performed for every strain on a single well plate (single colony), each replicate was measured at a range of inducer concentrations, and 3 replicate plates were measured with a different colony for each plate resulting in a total of 6 replicates at a given inducer concentration. Data analysis was performed by identifying the peak in the growth curve for each strain and each replicate (*X. autotrophicus* growth curves typically decrease after a peak at the end of exponential phase due to a change in cell shape). The peak OD_600_ was averaged over 3 datapoints, and the corresponding fluorescence data was averaged at the same 3 timepoints. The fluorescence was normalized to the optical density. In some cases, when the OD_600_ or fluorescence varied highly over the growth curve, exponential smoothing with α = 0.9 was used to identify the averaged OD_600_ or fluorescence intensity. The average fluorescence for all replicates at a given inducer concentration over a range of inducer concentrations is plotted in Fig. 5a–c and the error bars correspond to the standard deviation.

#### Termination Efficiencies

Cultures were grown for 17 h and the OD_600_, mRFP1 fluorescence (λ_ex_ = 577 nm; λ_em_ = 607 nm; gain 125), and GFP fluorescence (λ_ex_ = 485 nm; λ_em_ = 515 nm; gain 60) were measured every 15 min. 6 replicates were performed for every strain on a single well plate (single colony), and three replicate plates were measured with a different colony for each plate, giving a total of 18 replicates. Data analysis was performed by identifying the peak in the growth curve for each strain and replicate. The fluorescence value was averaged for the three RFP datapoints and three GFP fluorescence datapoints at the same timepoint as the growth peak. Termination efficiency was calculated by the following formula:

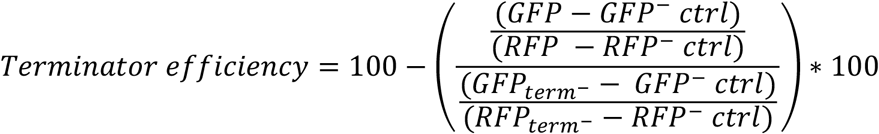

In each case, background green fluorescence was subtracted with a GFP^−^ control strain and background red fluorescence was subtracted with an RFP^−^ control strain. For a given strain, background-subtracted green fluorescence was divided by background-subtracted red fluorescence (numerator ratio). This same calculation was also applied to a terminator^−^ control (denominator ratio). The ratio of the two was then multiplied by 100, and this number was subtracted from 100 to give a termination efficiency with a % unit. Average termination efficiency for each terminator was plotted in Fig. 6, with the error bars corresponding to the standard deviations. Statistical analysis was performed by using a two-sample student t-test with unequal variances and determining whether the P value was > or ≤ 0.05.

#### Carbon Source Growth Experiments

50 mL cultures of the pJ23111 and pCin strains were grown overnight in YESMM, spun down, washed with fresh media, and resuspended in MM. The cells were used to inoculate fresh 3 mL cultures with either 0.2% methanol or 4 mM dichloroethane as the carbon source. 0.001% yeast extract was added as a vitamin mixture to facilitate the transition between carbon sources^17^ (this yeast extract concentration is insufficient as a carbon source; cultures with just 0.001% yeast extract do not grow). The cultures were sealed in tubes to prevent vaporization of the carbon source, and 200 uL of each culture were sampled at intervals to measure both the OD_600_ and the mRFP1 fluorescence (λ_ex_ = 577 nm; λ_em_ = 607 nm; gain 125).

## Author Contributions

R.S.S. conceived of the project and supervised the research; A.F.V. and R.S.S. designed the experiments, performed the research, and analyzed the data; A.F.V. and R.S.S wrote the manuscript.

## Data Deposition

The majority of the data is presented in this document and the supplementary information. All other data relevant to the manuscript is available upon request.

## Acknowledgements

R.S.S. is supported by the Burroughs Wellcome Fund grant (G-1021871.01) and startup funding through Northeastern University. The authors thank Prof. Benjamin Woolston and Prof. Edward Geisinger for plasmids and Prof. Ke Zhang for use of the real-time PCR instrument.

## References

(1) Adesina, O.; Anzai, I. A.; Avalos, J. L.; Barstow, B. Embracing Biological Solutions to the Sustainable Energy Challenge. Chem 2017, 2 (1), 20–51.

(2) Liao, J. C.; Mi, L.; Pontrelli, S.; Luo, S. Fuelling the Future: Microbial Engineering for the Production of Sustainable Biofuels. Nat. Rev. Microbiol. 2016, 14 (5), 288–304.

(3) Li, H.; Opgenorth, P. H.; Wernick, D. G.; Rogers, S.; Wu, T.-Y.; Higashide, W.; Malati, P.; Huo, Y.-X.; Cho, K. M.; Liao, J. C. Integrated Electromicrobial Conversion of CO2 to Higher Alcohols. Science 2012, 335 (6076), 1596–1596.

(4) Torella, J. P.; Gagliardi, C. J.; Chen, J. S.; Bediako, D. K.; Colón, B.; Way, J. C.; Silver, P. A.; Nocera, D. G. Efficient Solar-to-Fuels Production from a Hybrid Microbial–Water-Splitting Catalyst System. Proc. Natl. Acad. Sci. USA 2015, 112 (8), 2337–2342.

(5) Liu, C.; Colón, B. C.; Ziesack, M.; Silver, P. A.; Nocera, D. G. Water Splitting– Biosynthetic System with CO2 Reduction Efficiencies Exceeding Photosynthesis. Science 2016, 352 (6290), 1210–1213.

(6) Liu, C.; Sakimoto, K. K.; Colón, B. C.; Silver, P. A.; Nocera, D. G. Ambient Nitrogen Reduction Cycle Using a Hybrid Inorganic–Biological System. Proc. Natl. Acad. Sci. USA 2017, 114 (25), 6450–6455.

(7) Ryu, M.-H.; Zhang, J.; Toth, T.; Khokhani, D.; Geddes, B. A.; Mus, F.; Garcia-Costas, A.; Peters, J. W.; Poole, P. S.; Ané, J.-M.; Voigt, C. A. Control of Nitrogen Fixation in Bacteria That Associate with Cereals. Nat. Microbiol. 2019, 5, 314–330.

(8) Mus, F.; Crook, M. B.; Garcia, K.; Garcia Costas, A.; Geddes, B. A.; Kouri, E. D.; Paramasivan, P.; Ryu, M.-H.; Oldroyd, G. E. D.; Poole, P. S.; Udvardi, M. K.; Voigt, C. A.; Ané, J.-M.; Peters, J. W. Symbiotic Nitrogen Fixation and the Challenges to Its Extension to Nonlegumes. Appl. Environ. Microbiol. 2016, 82 (13), 3698–3710.

(9) Haskett, T. L.; Karunakaran, R.; Batista, M. B.; Dixon, R.; Poole, P. S. Control of Nitrogen Fixation and Ammonia Excretion in Azorhizobium Caulinodans. PLoS Genet. 2022, 18 (6), e1010276.

(10) Pankievicz, V. C. S.; Irving, T. B.; Maia, L. G. S.; Ané, J.-M. Are We There yet? The Long Walk towards the Development of Efficient Symbiotic Associations between Nitrogen-Fixing Bacteria and Non-Leguminous Crops. BMC Biol. 2019, 17 (1), 99.

(11) Kyriakou, V.; Garagounis, I.; Vourros, A.; Vasileiou, E.; Stoukides, M. An Electrochemical Haber-Bosch Process. Joule 2020, 4 (1), 142–158.

(12) Rylott, E. L.; Bruce, N. C. How Synthetic Biology Can Help Bioremediation. Curr. Opin. Chem. Biol. 2020, 58, 86–95.

(13) Janssen, D. B.; Pries, F.; van der Ploeg, J.; Kazemier, B.; Terpstra, P.; Witholt, B. Cloning of 1,2-Dichloroethane Degradation Genes of Xanthobacter Autotrophicus GJ10 and Expression and Sequencing of the dhlA Gene. J. Bacteriol. 1989, 171 (12), 6791–6799.

(14) Janssen, D. B.; Scheper, A.; Dijkhuizen, L.; Witholt, B. Degradation of Halogenated Aliphatic Compounds by Xanthobacter Autotrophicus GJ10. Appl. Environ. Microbiol. 1985, 49 (3), 673–677.

(15) Chen, Y.-J.; Leung, P. M.; Wood, J. L.; Bay, S. K.; Hugenholtz, P.; Kessler, A. J.; Shelley, G.; Waite, D. W.; Franks, A. E.; Cook, P. L. M.; Greening, C. Metabolic Flexibility Allows Bacterial Habitat Generalists to Become Dominant in a Frequently Disturbed Ecosystem. IMSE J. 2021, 15 (10), 2986–3004.

(16) Loos, D.; Filho, A. P. da C.; Dutilh, B. E.; Barber, A. E.; Panagiotou, G. A Global Survey of Host, Aquatic, and Soil Microbiomes Reveals Shared Abundance and Genomic Features between Bacterial and Fungal Generalists. Cell Rep. 2024, 43 (4).

(17) Wiegel, J. K. W. The Genus Xanthobacter. In The Prokaryotes Vol. 5: Proteobacteria: Alpha and Beta Subclasses; Brenner, D. J., Krieg, N. R., Garrity, G. M., Staley, J. T., Boone, D. R., Vos, P., Goodfellow, M., Rainey, F. A., Schleifer, K.-H., Eds.; Springer-Verlag: New York, 2006; Vol. 5, pp 290–314.

(18) Weaver, C. A.; Lidstrom, M. E. Methanol Dissimilation in Xanthobacter H4-14: Activities, Induction and Comparison to Pseudomonas AM1 and Paracoccus Denitrificans. Microbiol. 1985, 131 (9), 2183–2197.

(19) Weaver, C. A.; Lidstrom, M. E. Isolation, Complementation and Partial Characterization of Mutants of the Methanol Autotroph Xanthobacter H4-14 Defective in Methanol Dissimilation. Microbiol. 1987, 133 (7), 1721–1731.

(20) Small, F. J.; Ensign, S. A. Carbon Dioxide Fixation in the Metabolism of Propylene and Propylene Oxide by Xanthobacter Strain Py2. J. Bacteriol. 1995, 177 (21), 6170–6175.

(21) Allen, J. R.; Ensign, S. A. Carboxylation of Epoxides to Beta-Keto Acids in Cell Extracts of Xanthobacter Strain Py2. J. Bacteriol. 1996, 178 (5), 1469–1472.

(22) Zhou, N.-Y.; Chan Kwo Chion, C.K.; Leak, D. J. Cloning and Expression of the Genes Encoding the Propene Monooxygenase from Xanthobacter, Py2. Appl. Microbiol. Biotechnol. 1996, 44 (5), 582–588.

(23) Sluis, M. K.; Ensign, S. A. Purification and Characterization of Acetone Carboxylase from Xanthobacter Strain Py2. Proc. Natl. Acad. Sci. USA 1997, 94 (16), 8456–8461.

(24) Zhou, N.-Y.; Jenkins, A.; Chan Kwo Chion, C.K.N.; Leak, D. J. The Alkene Monooxygenase from Xanthobacter Strain Py2 Is Closely Related to Aromatic Monooxygenases and Catalyzes Aromatic Monohydroxylation of Benzene, Toluene, and Phenol. Appl. Environ. Microbiol. 1999, 65 (4), 1589–1595.

(25) Gogotov, J. N.; Schlegel, H. G. N2-Fixation by Chemoautotrophic Hydrogen Bacteria. Arch. Microbiol. 1974, 97, 359–362.

(26) Malik, K. A.; Schlegel, H. G. Chemolithoautotrophic Growth of Bacteria Able to Grow under N2-Fixing Conditions. FEMS Microbiol. Rev. 1981, 11, 63–67.

(27) De Bont, J. A. M.; Leijten, M. W. M. Nitrogen Fixation by Hydrogen-Utilizing Bacteria. Arch. Microbiol. 1976, 107 (3), 235–240.

(28) Berman-Frank, I.; Lundgren, P.; Chen, Y.-B.; Küpper, H.; Kolber, Z.; Bergman, B.; Falkowski, P. Segregation of Nitrogen Fixation and Oxygenic Photosynthesis in the Marine Cyanobacterium Trichodesmium. Science 2001, 294 (5546), 1534–1537.

(29) Gardner, J. J.; Hodge, B.-M. S.; Boyle, N. R. Investigating the Unique Ability of Trichodesmium To Fix Carbon and Nitrogen Simultaneously Using MiMoSA. mSystems 2023, 8 (1), e00601–20.

(30) Hu, X.; Kerckhof, F.-M.; Ghesquière, J.; Bernaerts, K.; Boeckx, P.; Clauwaert, P.; Boon, N. Microbial Protein out of Thin Air: Fixation of Nitrogen Gas by an Autotrophic Hydrogen-Oxidizing Bacterial Enrichment. Environ. Sci. Technol. 2020, 54 (6), 3609–3617.

(31) Sherbo, R. S.; Silver, P. A.; Nocera, D. G. Riboflavin Synthesis from Gaseous Nitrogen and Carbon Dioxide by a Hybrid Inorganic-Biological System. Proc. Natl. Acad. Sci. USA 2022, 119 (37), e2210538119.

(32) Kerckhof, F.-M.; Sakarika, M.; Van Giel, M.; Muys, M.; Vermeir, P.; De Vrieze, J.; Vlaeminck, S. E.; Rabaey, K.; Boon, N. From Biogas and Hydrogen to Microbial Protein Through Co-Cultivation of Methane and Hydrogen Oxidizing Bacteria. Front. Bioeng. Biotechnol. 2021, 9, 733753.

(33) Larsen, R. A.; Wilson, M. M.; Guss, A. M.; Metcalf, W. W. Genetic Analysis of Pigment Biosynthesis in Xanthobacter Autotrophicus Py2 Using a New, Highly Efficient Transposon Mutagenesis System That Is Functional in a Wide Variety of Bacteria. Arch. Microbiol. 2002, 178 (3), 193–201.

(34) Swaving, J.; van Leest, W.; van Ooyen, A. J. J.; de Bont, J. A. M. Electrotransformation of Xanthobacter Autotrophicus GJ10 and Other Xanthobacter Strains. J. Microbiol. Methods 1996, 25 (3), 343–348.

(35) Riley, L. A.; Guss, A. M. Approaches to Genetic Tool Development for Rapid Domestication of Non-Model Microorganisms. Biotechnol. Biofuels 2021, 14 (1), 30.

(36) Schuster, L. A.; Reisch, C. R. A Plasmid Toolbox for Controlled Gene Expression across the Proteobacteria. Nucleic Acids Res. 2021, 49 (12), 7189–7202.

(37) Cook, T. B.; Rand, J. M.; Nurani, W.; Courtney, D. K.; Liu, S. A.; Pfleger, B. F. Genetic Tools for Reliable Gene Expression and Recombineering in Pseudomonas Putida. J. Ind. Microbiol. Biotechnol. 2018, 45 (7), 517–527.

(38) Venkataraman, M.; Yñigez-Gutierrez, A.; Infante, V.; MacIntyre, A.; Fernandes-Júnior, P. I.; Ané, J.-M.; Pfleger, B. Synthetic Biology Toolbox for Nitrogen-Fixing Soil Microbes. ACS Synth. Biol. 2023, 12 (12), 3623–3634.

(39) Rondthaler, S. N.; Sarker, B.; Howitz, N.; Shah, I.; Andrews, L. B. Toolbox of Characterized Genetic Parts for Staphylococcus Aureus. ACS Synth. Biol. 2024, 13 (1), 103–118.

(40) Teh, M. Y.; Ooi, K. H.; Danny Teo, S. X.; Bin Mansoor, M. E.; Shaun Lim, W. Z.; Tan, M. H. An Expanded Synthetic Biology Toolkit for Gene Expression Control in Acetobacteraceae. ACS Synth. Biol. 2019, 8 (4), 708–723.

(41) Meyer, R. Replication and Conjugative Mobilization of Broad Host-Range IncQ Plasmids. Plasmid 2009, 62 (2), 57–70.

(42) Jain, A.; Srivastava, P. Broad Host Range Plasmids. FEMS Microbiol. Lett. 2013, 348 (2), 87–96.

(43) Caspi, R.; Pacek, M.; Consiglieri, G.; Helinski, D. R.; Toukdarian, A.; Konieczny, I. A Broad Host Range Replicon with Different Requirements for Replication Initiation in Three Bacterial Species. EMBO J. 2001, 20 (12), 3262–3271.

(44) Anindyajati Artarini, A.A.; Riani, C.; Retnoningrum, D. S. Plasmid Copy Number Determination by Quantitative Polymerase Chain Reaction. Sci. Pharm. 2016, 84 (1), 89– 101.

(45) Durland, R. H.; Toukdarian, A.; Fang, F.; Helinski, D. R. Mutations in the trfA Replication Gene of the Broad-Host-Range Plasmid RK2 Result in Elevated Plasmid Copy Numbers. J. Bacteriol. 1990, 172 (7), 3859–3867.

(46) Tait, R. C.; Close, T. J.; Rodriguez, R. L.; Kado, C. I. Isolation of the Origin of Replication of the IncW-Group Plasmid pSa. Gene 1982, 20 (1), 39–49.

(47) Anderson Promoter Collection, 2018. https://parts.igem.org/Promoters/Catalog/Anderson.

(48) Chen, Y.-J.; Liu, P.; Nielsen, A. A. K.; Brophy, J. A. N.; Clancy, K.; Peterson, T.; Voigt, C. A. Characterization of 582 Natural and Synthetic Terminators and Quantification of Their Design Constraints. Nat. Methods 2013, 10 (7), 659–664.

(49) Gale, G. A. R.; Wang, B.; McCormick, A. J. Evaluation and Comparison of the Efficiency of Transcription Terminators in Different Cyanobacterial Species. Front. Microbiol. 2021, 11.

