## Supplementary Information for "Creating a genetic toolbox for the carbon-fixing, nitrogen-fixing and dehalogenating bacterium *Xanthobacter autotrophicus*"

**Table S1.** Yeast extract succinate minimal media (YESMM) components.

| **Component** | **Concentration (g $L^{-1}$)** |
|---|---|
| Yeast extract | 0.01% w/v |
| Succinate | 0.1% w/v |
| $K_2HPO_4$ | 1.55 |
| $NaH_2PO_4$ | 0.85 |
| $NH_4Cl$ | 2 |
| $MgCl_2 \cdot 6H_2O$ | 0.035 |
| $(NH_4)_2SO_4$ | 0.1 |
| TMM | 0.2 mL $L^{-1}$ |
| **Trace Mineral Mixture (TMM)** | **Concentration (g 100 $mL^{-1}$)** |
| EDTA | 5 |
| $ZnSO_4 \cdot 7H_2O$ | 2.2 |
| $CaCl_2$ | 0.734 |
| $MnCl_2 \cdot 4H_2O$ | 0.323 |
| $FeSO_4 \cdot 7H_2O$ | 0.499 |
| $(NH_4)_6Mo_7O_{24} \cdot 4H_2O$ | 0.11 |
| $CuSO_4 \cdot 5H_2O$ | 0.157 |
| $CoCl_2 \cdot 6H_2O$ | 0.161 |
| KOH | adjust to pH 6 |

**Table S2.** List of all plasmids used in this study.

| **Plasmid Name** | **Plasmid Description** | **Part Tested** | **Source** |
|---|---|---|---|
| **pWLxR5** | Sa ori, gmR | Origin of replication | Schuster et al. *Nucleic Acids Res.*, 2021 addgene # 149490 |
| **pFCiR2** | RSF1010 ori; cinR pCin inducible promoter system | Origin of replication | Schuster et al. *Nucleic Acids Res.*, 2021 addgene #167512 |
| **pBADTrfp** | BBR1 ori | Origin of replication | Hillson Lab, addgene #99382 |
| **pFGM1** | gmR | N/A | Choi et al. *Nat Methods*, 2005 addgene #64949 |
| **pUC18** | ampR | N/A | Norrander et al. *Gene*, 1983 addgene #50004 |
| **pMMB207** | cmR | N/A | Morales et al. *Gene*, 1991 ATCC 37809 |
| **pRSS022** | gmR | Antibiotic resistance | This study |
| **pRSS023** | ampR | Antibiotic resistance | This study |

| | | | |
|---|---|---|---|
| **pRSS024** | cmR | Antibiotic resistance | This study |
| **pRSS000** | pRpsM rfp; RK2 ori | Constitutive promoter | This study |
| **pRSS011** | pLacIq | Constitutive promoter | This study |
| **pRSS018** | pJ23102 rfp | Constitutive promoter | This study |
| **pRSS019** | pJ23111 rfp | Constitutive promoter | This study |
| **pRSS020** | pJ23107 rfp | Constitutive promoter | This study |
| **pRSS021** | pJ23103 rfp | Constitutive promoter | This study |
| **pRSS030** | pRpsM control ($RFP^{-}$) | Constitutive promoter | This study |
| **pBbS2k** | tetR pTet inducible promoter system | N/A | Lee et al. *J. Biol. Eng.,* 2011 addgene #35330 |
| **pBba5c** | lacI pLac inducible promoter system, cmR | N/A | Lee et al. *J. Biol. Eng.*, 2011 addgene #35281 |
| **pRK2-AraE** | araE arabinose importer, gmR | N/A | Cook et al. *J. Ind. Microbiol. Biotechnol.* 2018 addgene #110141 |
| **pRSS008** | araC pBAD rfp | Inducible promoter – arabinose | This study |
| **pRSS010** | cinR pCin rfp | Inducible promoter – homoserine lactone | This study |
| **pRSS031** | cinR pCin control ($RFP^{-}$) | Inducible promoter – homoserine lactone control | This study |
| **pRSS035** | araE araC pBAD rfp | Inducible promoter – arabinose with arabinose importer | This study |
| **pRSS037** | lacI pLac rfp | Inducible promoter – lactose/IPTG | This study |
| **pRSS038** | tetR pTet rfp | Inducible promoter – tetracycline | This study |
| **pFAB803** | rfp T7TE term gfp | N/A | Mutalik et al. Nat. Methods, 2013 addgene #47847 |
| **pFAB808** | rfp rrnB term gfp | N/A | Mutalik et al. Nat. Methods, 2013 addgene #47853 |
| **pFAB815** | rfp rnpB term gfp | N/A | Mutalik et al. Nat. Methods, 2013 addgene #47854 |
| **pFAB822** | rfp B1006 term gfp | N/A | Mutalik et al. Nat. Methods, 2013 addgene #47856 |
| **pRSS012** | rfp rrnB term gfp | Terminator | This study |

| | | | |
|---|---|---|---|
| **pRSS013** | rfp B1006 term gfp | Terminator | This study |
| **pRSS014** | rfp T7TE term gfp | Terminator | This study |
| **pRSS015** | rfp rnpB term gfp | Terminator | This study |
| **pRSS016** | rfp B0015 term gfp | Terminator | This study |
| **pRSS028** | rfp no term gfp (term$^{-}$) | Terminator | This study |
| **pRSS029** | rfp B0015 term (GFP$^{-}$) | Terminator | This study |

*Unless otherwise noted, all plasmids have an RK2 origin of replication, *kanR* antibiotic resistance gene, an *mrfp1* reporter gene and a B0015 double terminator.

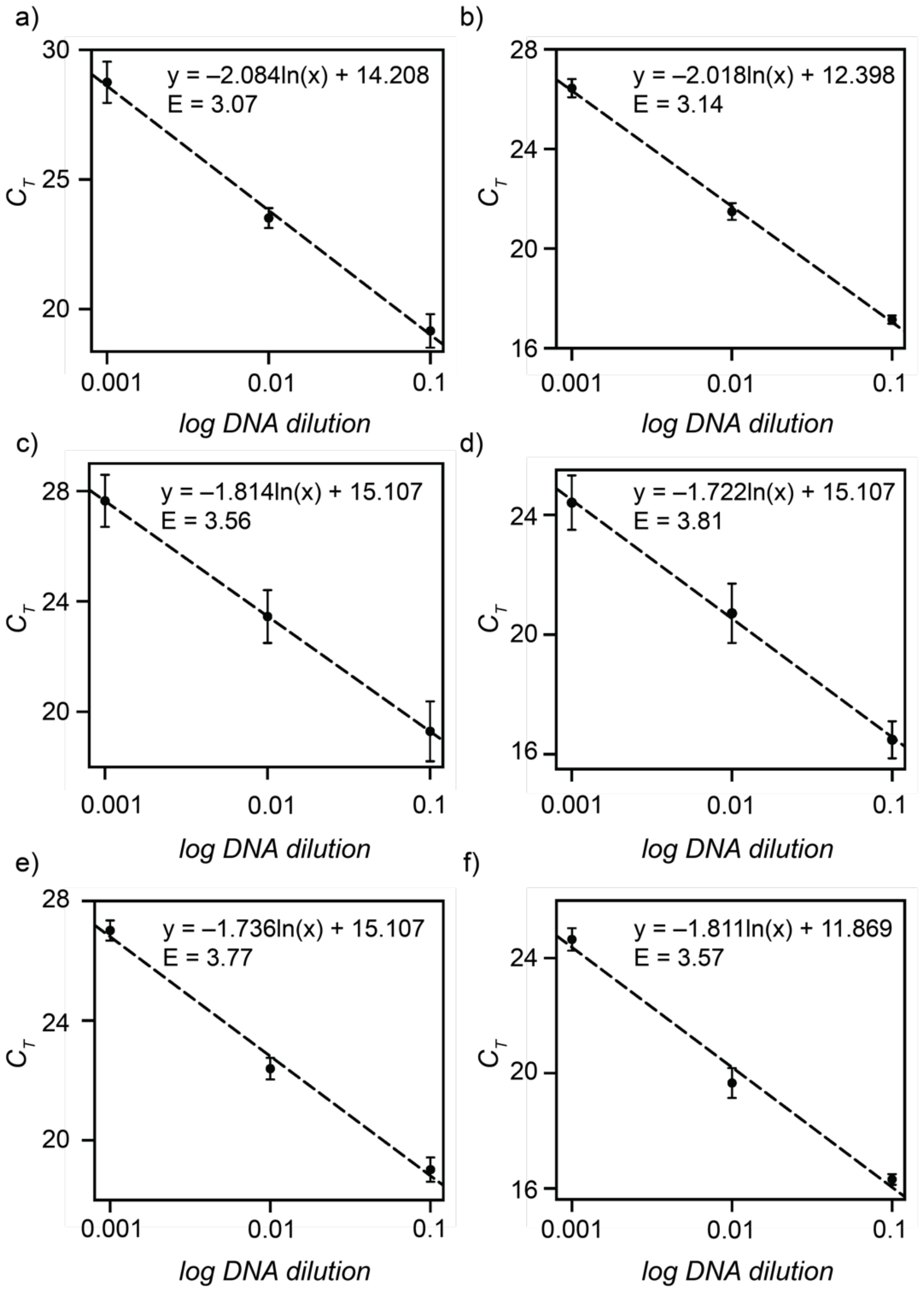


**Figure S1.** qPCR primer efficiencies for a, c, e) the chromosomal primer pair and b, d, f) the plasmid primer pair for RK2 (a, b), pSa (c, d) and BBR1 (e, f) origins of replication.

**Table S3.** Antibiotic concentrations for use in *X. autotrophicus*.

| Antibiotic | Concentration |
|---|---|
| kanamycin | 30 µg $mL^{-1}$ |
| gentamycin | 7.5 µg $mL^{-1}$ |
| ampicillin | 50 µg $mL^{-1}$ |
| chloramphenicol | 10 µg $mL^{-1}$ |

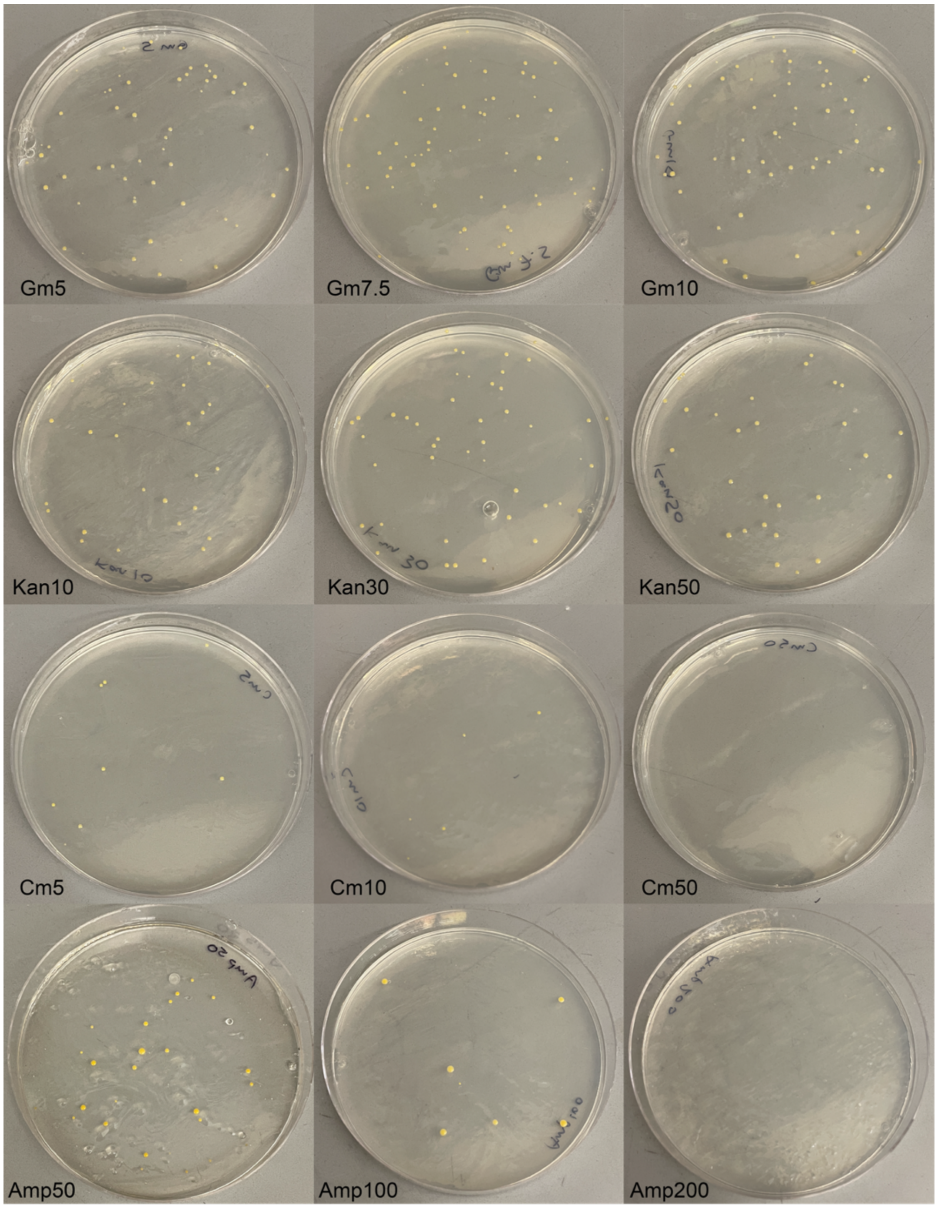


**Figure S2.** *X. autotrophicus* antibiotic resistant strain electrotransformation plating onto various antibiotics and antibiotic concentrations.

**Table S4.** Colony-forming units in antibiotic-containing cultures plated on antibiotic-containing plates.

| Antibiotic | Colony-forming Units |
|---|---|
| kanamycin | $1.15\times10^{8}$ cfu mL$^{-1}$ |
| gentamycin | $2.15\times10^{8}$ cfu mL$^{-1}$ |
| ampicillin | $3.05\times10^{7}$ cfu mL$^{-1}$ |
| chloramphenicol | $1.05\times10^{8}$ cfu mL$^{-1}$ |

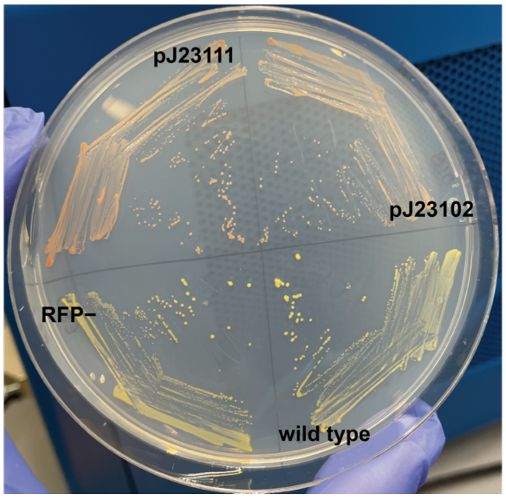


**Figure S3.** pJ23111 and pJ23102 constitutive promoters result in pink/orange colonies from high red fluorescence levels compared to the yellow empty vector (RFP$^{-}$) and wild type strains.

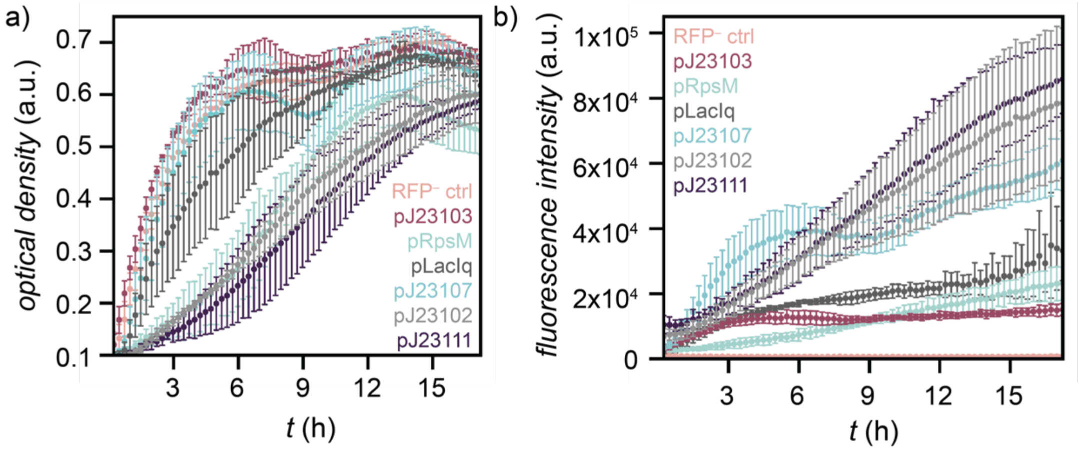


**Figure S4.** a) Optical density at 600 nm (growth) and b) red fluorescence intensity ($\lambda_{ex}$ = 577 nm; $\lambda_{em}$ = 607 nm) of the constitutive promoter strains.

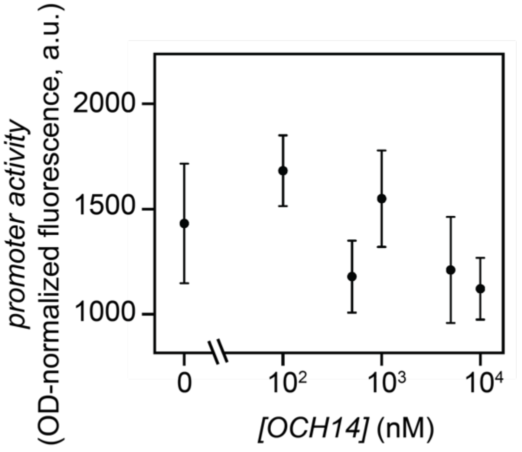


**Fig. S5.** Induction curve for a pCin inducible promoter empty vector control (RFP$^{-}$) to demonstrate a lack of fluorescence without the red fluorescent protein reporter.

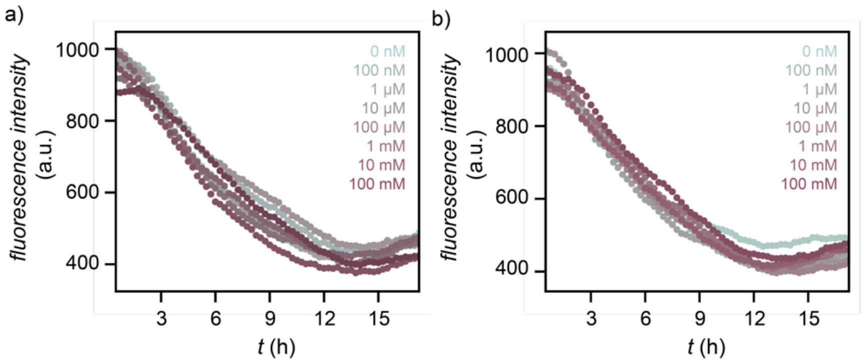


**Fig. S6.** Red fluorescence intensity ($\lambda_{ex}$ = 577 nm; $\lambda_{em}$ = 607 nm) of a) the pBAD arabinose inducible promoter and b) the pBAD inducible promoter with the AraE arabinose importer at various concentrations of arabinose inducer. The fluorescence is similar with no inducer and at every tested inducer concentration.

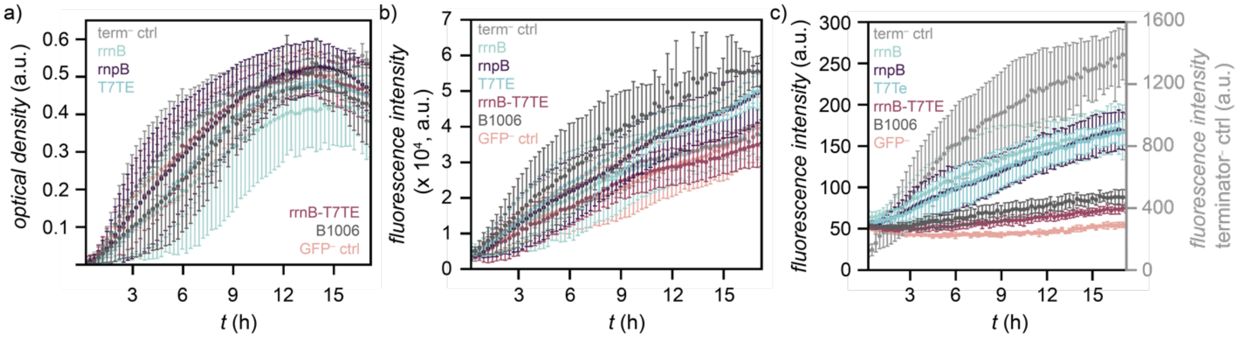


**Figure S7.** a) Optical density at 600 nm (growth), b) red fluorescence intensity ($\lambda_{ex}$ = 577 nm; $\lambda_{em}$ = 607 nm) and c) green fluorescence intensity ($\lambda_{ex}$ = 577 nm; $\lambda_{em}$ = 607 nm) of the terminator strains. All green fluorescence intensity is associated with the left y-axis except the terminator⁻ control which is associated with the right y-axis.
